# New insights into the Germline Genes and CDR3 Repertoire of the TCRβ chain in *Chiroptera*

**DOI:** 10.1101/2022.12.05.519110

**Authors:** Hao Zhou, Jun Li, Dewei Zhou, Yingjie Wu, Xingliang Wang, Jiang Zhou, Qingqing Ma, Xinsheng Yao, Long Ma

## Abstract

Bats are known to be natural reservoirs of many viruses, yet their unique immune system enables them to coexist with viruses without frequently exhibiting disease symptoms. The current understanding of the bat adaptive immune system is limited, as there is no database or tool capable of processing T-cell receptor (TCR) sequences for bats, and the diverse nature of the Chiroptera. We established a reference library of TCR-β germline genes by annotating three Chiroptera: The Greater Horseshoe Bat (Rhinolophus ferrumequinum, R. ferrumequinum), The Pale spear-nosed Bat (Phyllostomus discolor, P. discolor), and the Common Pipistrelle (Pipistrellus pipistrellus, P. pipistrellus). The distinct variations in the distribution of TRBV genes among the three types of bats could have a direct impact on the diversity of the TCR repertoire, as evidenced by the presence of conserved amino acids that indicate the T-cell recognition of antigens in bats is MHC-restricted. Furthermore, we conducted an analysis of the TCR-β repertoire in the Intermediate Horseshoe Bat (Rhinolophus affinis, R. affinis) using high-throughput sequencing (HTS). The bats’ TCR-β repertoire is formed through rearrangement of the V-D-J-C genes, with D-J/V-D deletions and insertion resulting a high diversity.

## 1 Introduction

Bats belong to the order Chiroptera, are the second largest order of mammals in the world (*1*). Despite carrying numerous virulent zoonotic viruses, bats do not often show serious clinical symptoms (*2–5*). This has led to increased interest in the differences between the bat immune system and those of other mammals.

Studies have identified several differences between the bat and human or mouse innate immune systems. For example, activation of patterns recognition receptors (PRRs) by RNA viruses, danger signals, or intracellular double-stranded DNA in humans or mice triggers the activation of NLR-family pyrin domain containing 3 (NLRP3) or Interleukin-1β (IL-1β) resulting in inflammation. In contrast, bats inhibit the transcriptional initiation of NLRP3, leading to reduced functionality of Interferon-inducible proteins AIM2 and IFI-16(*6–9*), as well as lower caspase-1 activity and IL-1β cleavage, resulting in overall reduced inflammation. A recent study found that deletion of the 358th serine site of the bat STING protein inhibits IFN secretion(*10*). Additionally, Pavlovich et al. reported a “high amplification and non-classical distribution of genes” at MHC-I loci in bats, suggesting that the combination of these abundant non-classically distributed MHC-I type genes with highly expressed NKG2, which contain inhibitory interaction motifs, increases the activation threshold of NK cells and reduces the response (*11, 12*).

The understanding of the adaptive immune system of bats is hindered by the lack of appropriate reagents and models for cellular biology studies, as well as the challenge of isolating viruses and the diverse nature of bat species (which belong to 2 orders, 21 families, and include more than 1350 species) (*13*).With the advancement of Next Generation Sequencing (NGS) technology, high-throughput sequencing (HTS) has become a commonly used method for analyzing the genome of species. The Bat1k Project and Vertebrate Genome Project (VGP) of high-quality bat genomic data has greatly advanced our understanding of the bat immune system, especially their response to viral infections(*14, 15*). Despite this, the annotation of the T/B cell receptor germline genes in bats is still limited. The TCR/BCR repertoire is a collection of all functional T or B cells in an individual’s circulatory system, each with its own antigen-specific receptor. Deep sequencing of the repertoire is widely used to study the adaptive immune system, assess an individual’s health status, develop antibodies, and detect and threat targeted diseases (*16*). Various tools, such as IMGT/HighV-QUEST, TRUST, MiXCR, have been used for processing TCR/BCR sequences (*17–19*). Our previous work involved annotation of R. ferrumequinum TRB loci, suggesting that the bat’s adaptive immune response is similar to humans and mice and that its TCR is rearranged from germline genes with high diversity(*20*). In this study, we extended the annotation to include the TRB loci of the P. discolor and P. pipistrellus, and compared the TRB loci of all three bats species. The TRBC Exon1 sequences of various bats were aligned, and primers were developed to amplify the TCR β-chain CDR3 repertoire of R. affinis using the 5’ Rapid Amplification of cDNA Ends (5’RACE) technique. Our study provides a crucial theoretical foundation, a novel research method, and a comparable database for studying genetic evolution and adaptive immune response in *Chiroptera.*

## 2 Materials and methods

### 1. TRB locus location and annotation

Figure 1A illustrates the experimental flow of this study. We located the TRB loci in the genome by identifying two genes, Monooxygenase DBH-like 2 (MOXD2) and EPHB receptor6 (EPHB6), which were positioned on the boundary of the TRB loci. There are two gene annotation methods for the regions between two specific genes: 1. IMGT-LIGMotif (*21*) and 12/23 RSS (Recombination Signal Sequence) scanning(*22*). For the IMGT-LIGMotif method, we selected representative animals from three different subjects: human(primate), mouse(rodents), and pig(artiodactyla) for the search. To make sure that no information was missed due to evolutionary divergence, we obtained the TRBV genes of all known species in IMGT, which only allows retrieval of genes for one species at a time. We then used Geneious Prime (Version 2022.2.1) to map these genes to the TRB loci of the three bat species. The 12/23 RSS scanning method was similar to previous study(*20*). Briefly, we screened all RSS motifs within the three bat TRB loci and searched for V/D/J genes upstream and downstream of the RSS. The annotation result, including the number of genes and locus composition information, are summarized in Table 1. Using Easyfig (Version 2.2.5), we compared the homology of the TRB loci among the three annotated bats with a threshold of 70% identity. Following the IMGT guidelines(*23*), we labeled the key components of each germline gene within the locus after annotation with Geneious Prime and exported the annotation file in Genbank format(Supplementary Data1 and Supplement Figure 1).

**Figure 1.**
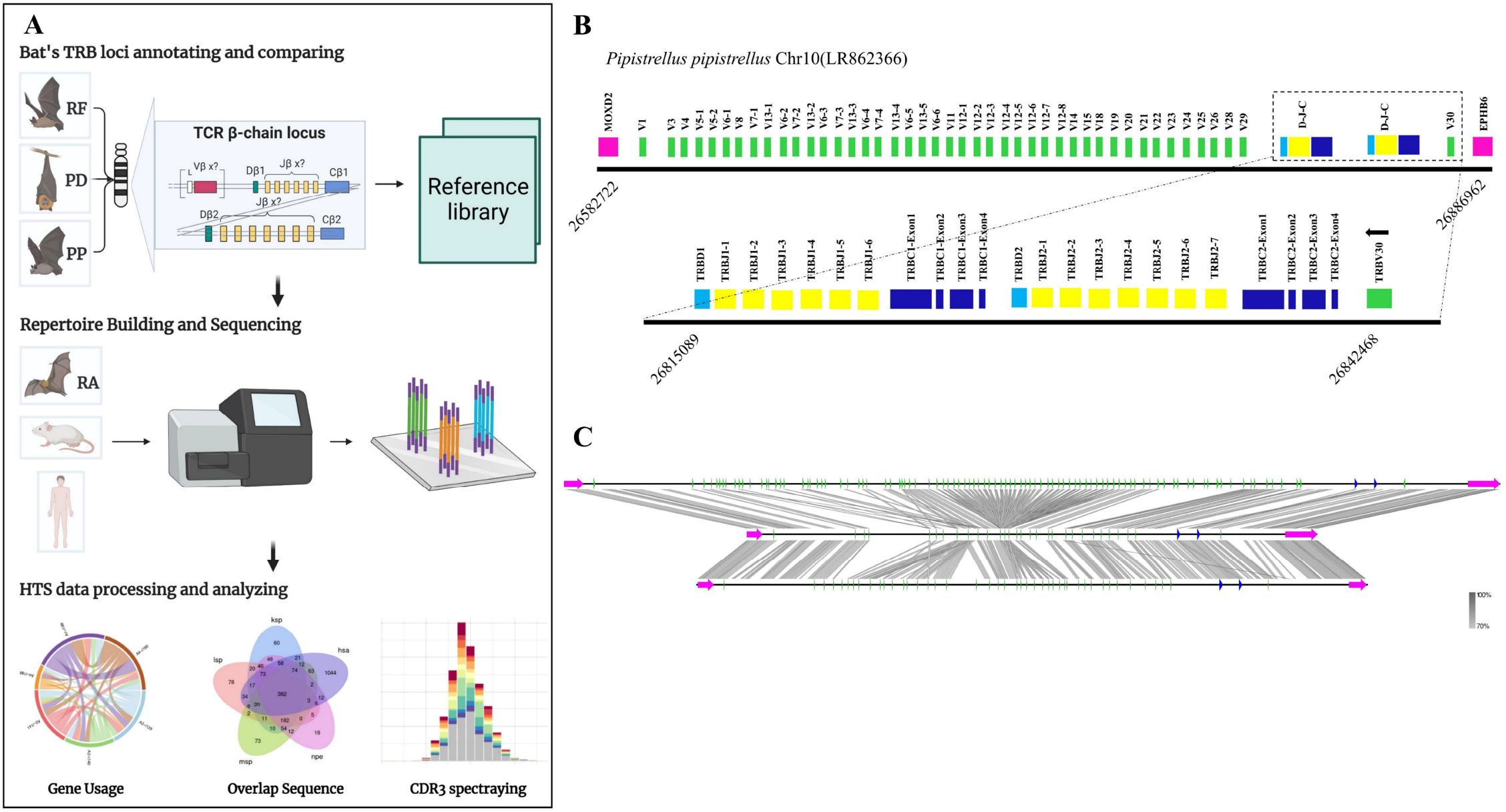
The TRB loci annotation and correlation analysis of three bat species. (A) Overview of this study design. (B) P. pipistrellus TRB loci on Chromosome 10, with colored boxes representing each annotated gene, including 44 TRBV genes, 2 TRBD genes, 13 TRBJ genes, and 2 TRBC genes. (C) A genome homology comparison map of the TRB loci for the three annotated bat species.

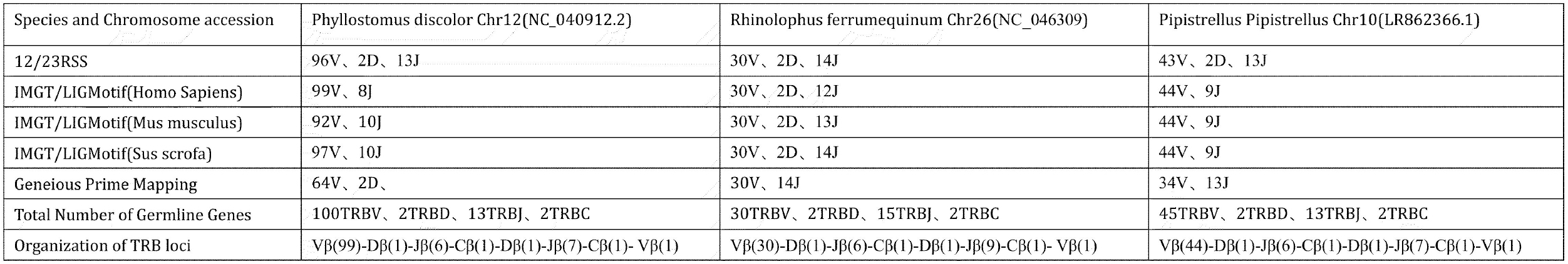

### 2. Naming of Germline Genes and Clustering of TRBV families

The naming of each germline gene was done according to the IMGT guidelines. For V genes, if the nucleotide identity was more than 75%, it was considered as belonging to the same family. Using MEGA7 (version 7.0.26), we constructed a phylogenetic tree of TRBV nucleotide sequences employing the Neighbor-Joining method to verify familial homology. The TRBV genes from human, mouse, and pig were obtained from IMGT-GeneDB (https://www.imgt.org/genedb/), with the accession numbers listed in the Supplementary Table 1. The criteria for selecting genes were: (1) only functional genes and ORFs, and (2) one gene per family. The distribution of TRBV gene families in twelve species belonging to four different mammalian subjects was compared to understand the evolutionary direction of the V gene, including Artiodactyla (cattle, sheep, and pigs), Carnivores (dogs, cats, and ferrets), and Primates (humans, crab-eating monkeys, and rhesus monkeys). The D, J, and C genes were named based on their position within the cluster they belong to in the locus.

### 3. Establishment of Reference Library

To establish a reference library, we aligned the amino acid sequences of the germline V and J genes using Geneious Prime (Clustal Omega). We gave priority alignment to three conserved sites - 23CYS, 41TRP, and 104CYS - based on the known structure and specific positions of mammalian TRBV (*23*). The structural domains were classified according to the results of the alignment: V genes included FR1 (1-26), CDR1 (27-38), FR2 (39-55), CDR2 (56-65), FR3 (66-104), and CDR3 (105-117), while J genes included FR4 (phenylalanine to the end of the FGNG motif sequence). The functional descriptions of the germline genes were based on IMGT, taking into account stop codons, RSS, conserved amino acids, and splice sites. Information on pseudogenes and ORFs is provided in the Supplementary Table 2. The germline gene nucleic sequences and their structural domain information were exported as reference library files identifiable by MiXCR, to process bat TRB data.

### 4. Sample collection

Bat samples were collected in Xishui County, Zunyi City, Guizhou Province with the assistance from Zhou’s group. We used muscle tissues to identify their genetic information by amplifying the Cytb gene with designed primers (Supplementary Table 5). The PCR products were sequenced to determine the bats’ genotype based on E-value and Percent Identity. As we did not find any annotated bat species based on the BLAST results, we selected three R. affinis as our experimental model, as it was expected to have high homology with the target species R. ferrumequinum. In addition, spleen tissues from three 5-month-old BALB/c mice were used for RNA extraction and TCRβ chain library construction. Furthermore, we collected molecular cells from 5 ml of peripheral blood from three individuals at the Affiliated Hospital of Zunyi Medical University. The Animal Protection and Ethics Committee of Zunyi Medical University approved the study, and work with humans was approved under permit number (2021)1-022. The bat and mouse project were approved under permit number (2018)2-261.

### 5. TCR-β chain repertoire sequencing and analysis

To construct and amplify the TCR-β library for R. affinis, we collected and aligned fifteen known bats’ TRBC Exon1 nucleotide sequences and designed primers in conserved region (accession number in Supplementary Table1). We used the 5’RACE method to prepare repertoires of R. affinis, with library construction and sequencing performed by ImmuQuad Company in Hangzhou, China. For species repertoire comparison, we used samples from humans and mice. We constructed the TCR-β repertoires of mice using spleen tissue with 5’RACE method, while for humans, we collected peripheral blood mononuclear cells (PBMC) from three individuals and used the multiplex PCR method for TCR-β repertoire construction. To analyze the sequencing results of bats, humans, and mice, we used MiXCR software (Version 3.0.13). Speciafically, we used MiXCR to assemble and present the sequencing results of bats, based on the library constructed from annotated genes. Table 2 and Supplementary Table 6 show the output data of the nine samples. To ensure the quality of repertoire, we examined the sequences of the bat1 sample, which contained 25 TRBV genes and 14 TRBJ genes (Figure 4E). The evolutionary relationship between the sequenced samples and the three annotated bats was determined by sequence attribution, Cytb, and TRBC gene alignment. To assess the homogeneity and heterogeneity of TCR-β repertoires among the three species, we used Productive-Clonetype sequences for subsequent analyses to exclude the effects of repertoire construction and sequencing, as well as physiological health status. Our analyses included: (1) statistics on repertoire overlap index among the three species; (2) usage of TRBV gene and TRBJ gene, V-J gene pairing, and tracking of conserved motifs; and (3) evaluation of CDR3 length and amino acid usage, as well as nucleotide deletion and insertion.

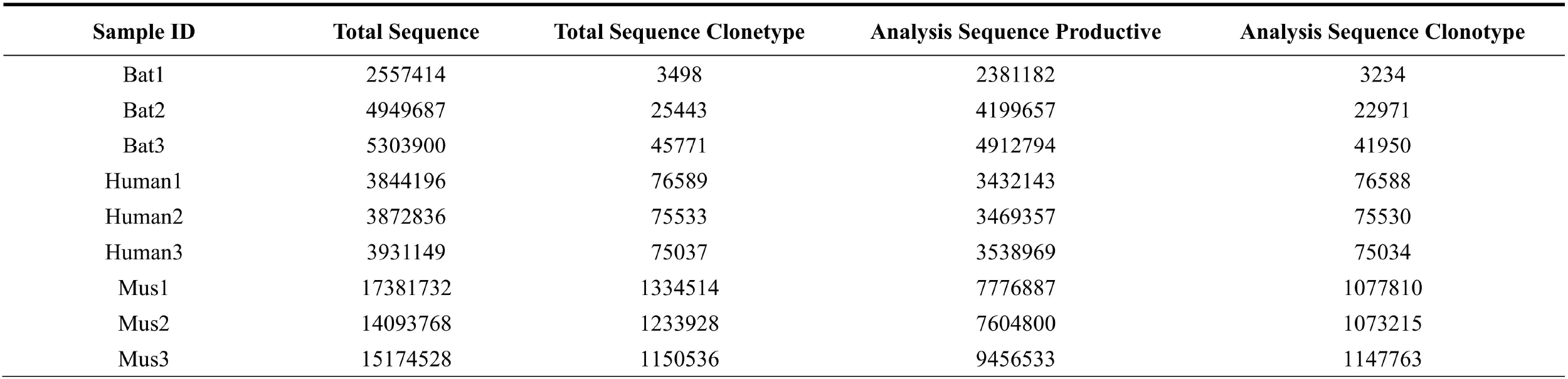

### 6. Statistical analysis

The figures were draw with GraphPad Prism (Version 8.0.2), R package “ggplot2”, and R package “Immunarch”. Data analysis was performed by R studio (v3.3.3) and GraphPad Prism. P-values were calculated with the aid of the t test. P<0.05 was considered statistically significant.

## 3 Results

### 1. TR loci annotation and Reference directory establishment

Table 1 records the summary of TRB loci using homologous genes and RSS methods. The TRB location in P. discolor was found on chromosome 10(NC_040912: 85517683-85943353), R. ferrumequinum on chromosome 26(NC_046309: 7911840-8171343), and P. pipstrellus on chromosome10(LR862366.1: 26582722-26886962). The TRB loci in the three bat species followed the classical mammalian structure, with differences in length distribution and orientation (Figure 1B, Supplementary Figure1). We identified 100 TRBV genes in P. discolor (including 2 pseudogenes), 30 in R. ferrumequinum, and 45 in P. pipistrellus, and a phylogenetic tree using the nucleotide sequences of TRBV genes reveal that all genes could be homologous to each other or to those found in pigs and humans, and no functional V gene families clustered separately in bats (Figure 2A). The main differences in the number of germline genes arose from internal duplications and deletions of the V gene family, with Chiroptera and Artiodactyla being more similar in their evolutionary direction (Figure 2B). Most of the TRBV and TRBJ gene families of the three bat species were more than 70% identity, except for the TRBV17 family (Figure 2C). A particular TRBV12 family of P. discolor containing 37 members was identified, which was close to almost all TRBV genes of P. pipistrellus and even more than the total TRBV genes of R. ferrumequinum.

**Figure2.**
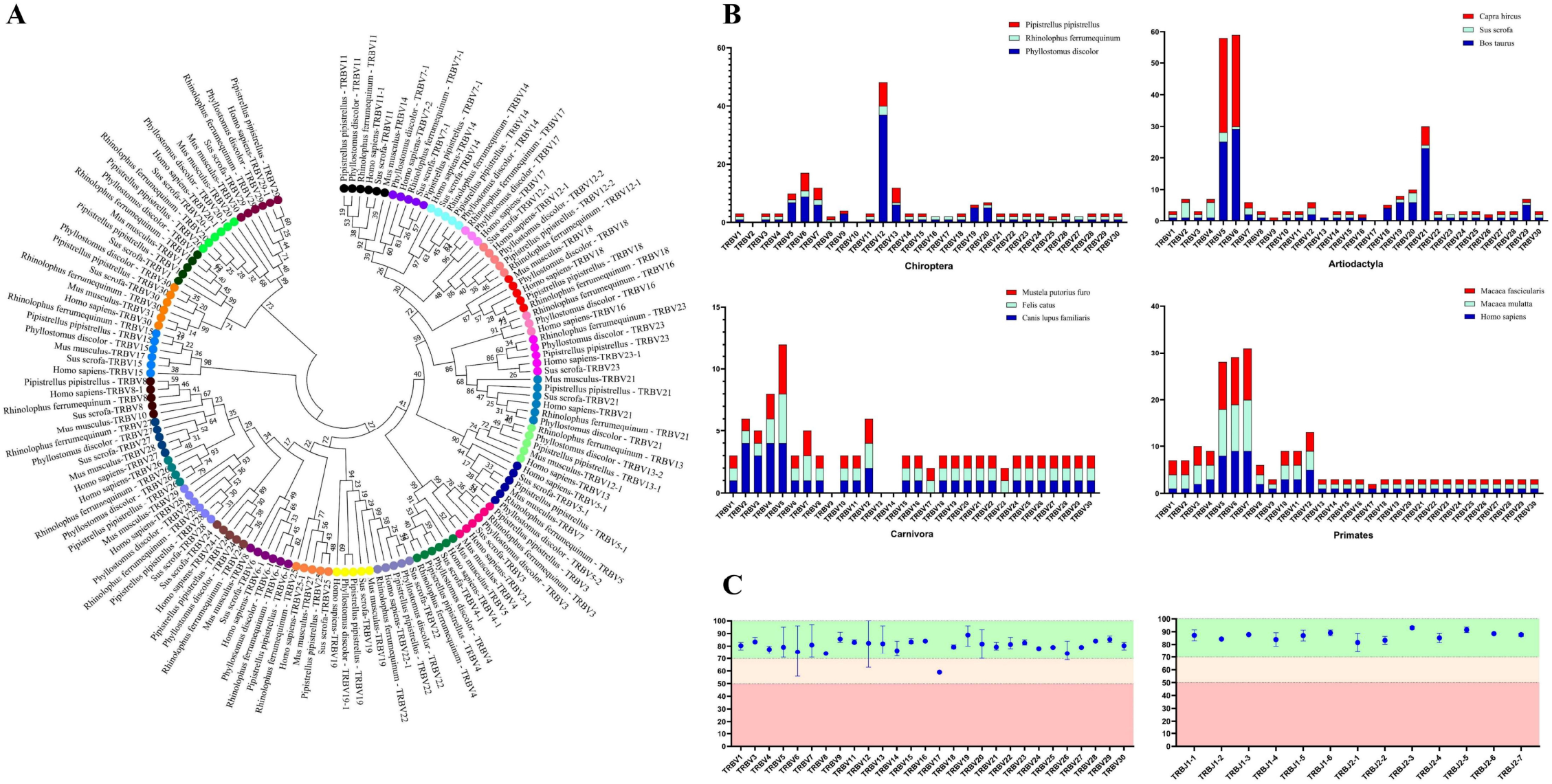
Analysis of TRBV gene homology in different bat species, as well as in human, mice, and pigs. (A) An NJ-phylogenetic tree was constructed from functional TRBV gene nucleotide sequences, with each color representing one family and bootstrap is 1000. (B) A comparison of TRBV gene family distribution in three mammalian orders (Carnivora, Artiodactyla, and Primates) with the three bat species is shown, with data from https://www.imgt.org/IMGTrepertoire/. (C) A comparison of the nucleotide sequence identity of TRBV and TRBJ genes is presented for the three annotated bat species.

In the Multiple Sequence Alignment (MSA) of TRBV genes, we identified several conserved sites, including 23Cys, 41Trp, 89Leu, and 104Cys, as well as Gln at position 6 and Tyr at position 42. Gln at position 44 was also highly conserved in P. pipistrellus. Additionally, most TRBV genes contained CASS motifs at the end (Figure 3A, Supplementary Figure 2A). A higher number of germline genes and more gene duplications in P. discolor resulted in a greater number of pseudogenes, 17%, compared to R. ferrumequinum (7%) and P. pipistrellus (11%). However, the number of functional genes (V and J genes) in P. discolor, which exceeded that of R. ferrumequinum and P. pipistrellus, was 97 (Supplementary Table 2). The Chiroptera, like primates and carnivores, contains two D-J-C clusters named according to their chromosomal location. The only difference between the three bat species in terms of the number of genes in each cluster was the number of J genes in the second cluster. All TRBJ genes in bats contain conserved FGNG motifs except for TRBJ2-3 in R. ferrumequinum (Figure 3B, Supplementary Figure 2B). In addition, all bats’ TRBD genes are guanine-rich sequences, with 23RSS and 12RSS downstream and upstream (Figure 3C). There were no significant differences in the RSS between bats and other mammals, and the most conserved site was the first four sites (CACA) of the heptamer, with three continuous adenosines in the middle position of the nonamer (Figure 3D, Supplementary Figure 3). We created a MiXCR reference library for HTS data analysis using nucleotide sequences and amino acid sequence MSA results. To organize the germline genes family of three bat species, we combined them into a single family with allelic nomenclature methods. (Supplementary Table 3).

**Figure3.**
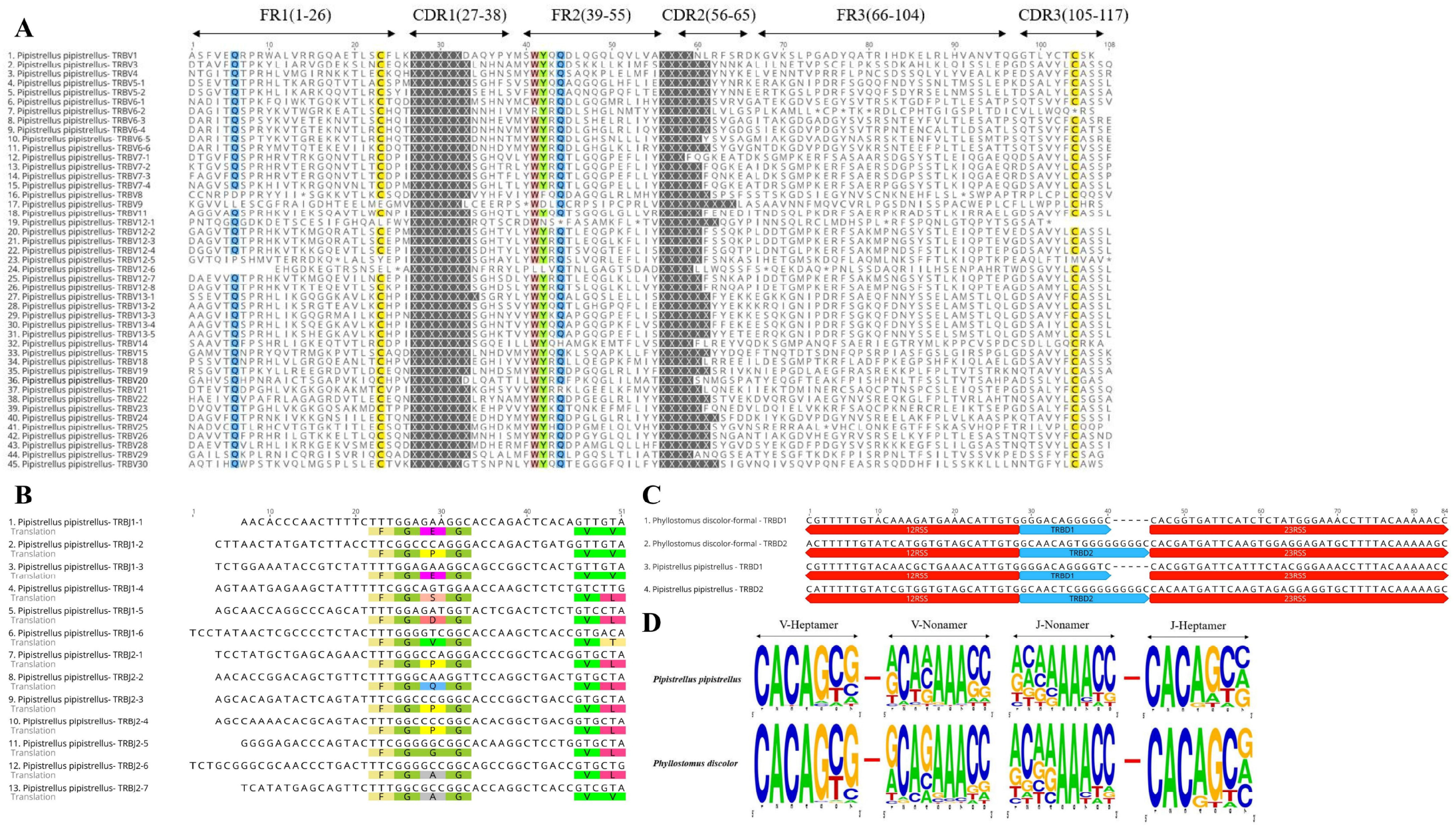
Annotated Germline Genes Display. (A) Displays 45 TRBV amino acid sequences with a consistency threshold of 90%. (B) Displays the nucleotide and amino acid sequences of 13 TRBJ genes with translated conserved FGNG motifs. (C) Shows four TRBD genes along with their 23RSS and 12RSS nucleotide sequences. (D) Compares the conserved sites of TRBV23RSS and TRBJ?2RSS in P. discolor and P. pipistrellus.

### 2. TCRβ Repertoire Constructing, Sequencing, Analyzing

The Cytb sequencing results of the three experimental bats were analyzed, revealing an average length of 488 bp and 98.5% identity. These bats were identified as R. affinis through BLAST analysis (Supplementary Table 4). Furthermore, the TCR) chain repertoire was constructed using 5’RACE by comparing the conserved regions of the TRBC Exon1 nucleotide sequences of fifteen types of bats and designing primers. The nucleotide sequence identity of 19 TRBC Exon1 ranged from 77.66% to 98.9%, with many fully conserved regions suitable for repertoire construction and sequencing (Figure 4A).

**Figure4.**
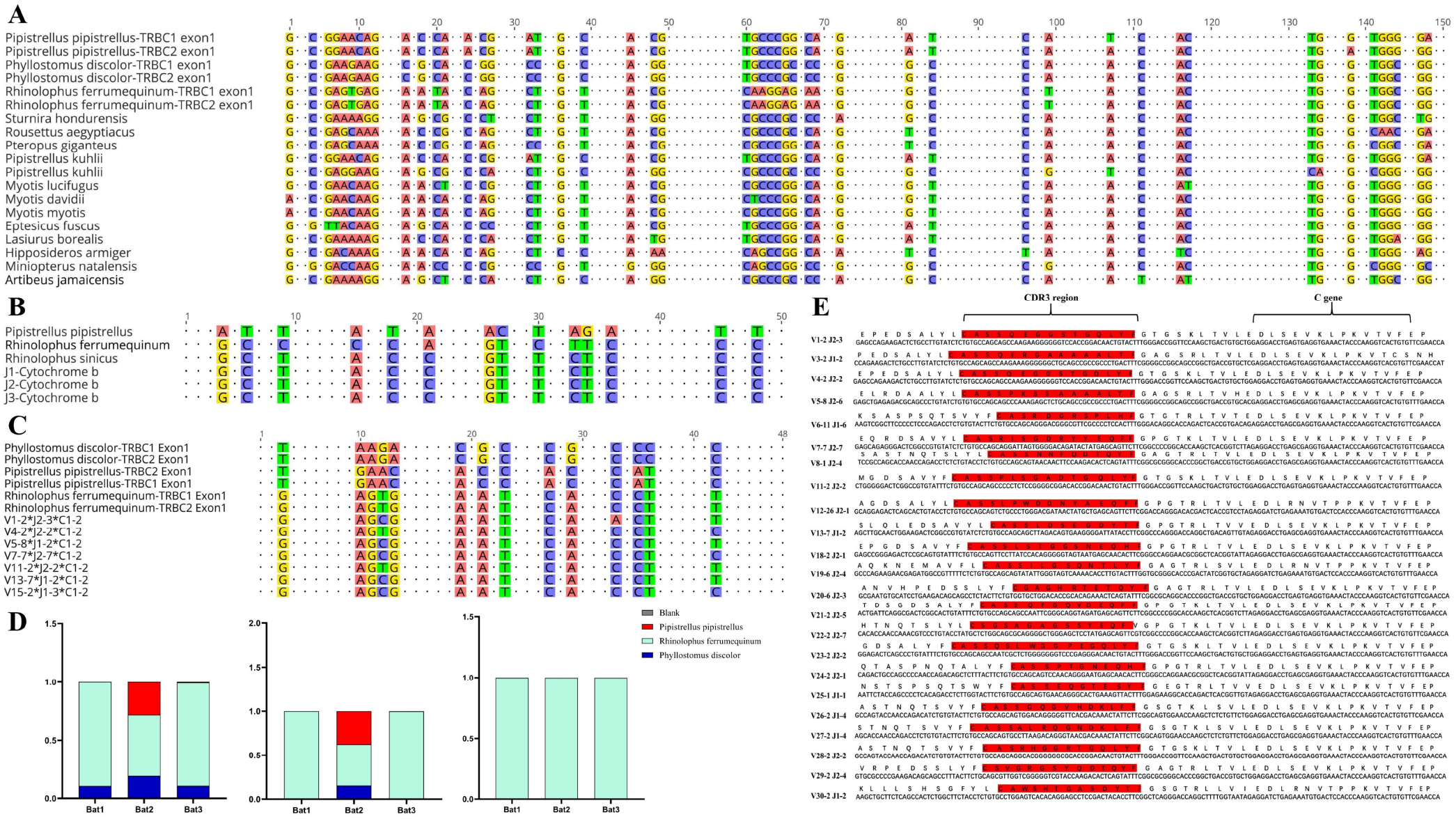
Homology analysis of R. affinis samples and annotated bat species. (A) Alignment of TRBC Exonl nucleotide sequences of 15 Chiroptera species used for primer design in the 5’RACE method of R. affinis. (B) Comparison of Cytb nucleotide sequence results of R. affinis samples with R discolor (PD), R. ferrumequinum (RF), and P. pipistrellus (PP). (C) Comparison of a partial C gene fragment from the batl sample with those of the three annotated bat species. (D) Attribution statistics of the sequencing results of the three R. affinis samples in the three annotated bat species. (E) Output of 25 TRBV genes and 14 TRBJ genes in the batl sample.

The sequencing was conducted by ImmuQuad Company and the library was confirmed by a peak between 600bp and 700bp in the R. affinis samples (Supplementary Figure 4). Raw data from nine samples were then processed by MiXCR for further analysis and can be accessed at NCBI_PRJNA877449 (SRX17468175, SRX17468176, and SRX17468177). The 5’race method yielded an average of 4.27 million TCR reads and 24,904 clonotypes per bat sample, 15 million TCR reads and 1.23 million clonotypes per mouse sample. The multiplex-PCR method resulted in 3.8 million TCR reads and 75,000 clonotypes per human sample. All samples, despite variations in sequencing depth across bat samples, met the CDR3 sequence depth analysis requirements for HTS sequencing, as indicated in Table 2.

For the sequencing results of R. affinis, to explore the degree of bat species divergence, we firstly compared the Cytb nucleotide sequence of the three experimental bats with R. ferrumequinum and P. pipistrellus (Figure 4B). The R. ferrumequinum (87.9%) had higher identity to R. affinis than P. pipistrellus (74.6%). Besides, the TRBC nucleotide fragment demonstrated identity between R. affinis with P. discolor, R. ferrumequinum and P. pipistrellus: 77.1%, 93.8%, 75.0% respectively (Figure 4C). After categorizing the sequencing results of the experimental samples R. affinis, we found that most of the sequences of R. affinis output sequences belonged to R. ferrumequinum (Figure 4D). The comparison results further indicated that the sequences of R. affinis were more similar to those of R. ferrumequinum. Then, we demonstrated the completeness of the repertoire by exporting 25 V and 14 J gene sequences from Bat1 samples (Figure 4E).

### 3. Comparison of CDR3 Repertoire in Bats, Humans, and Mice

Four indices (overlap coefficient, Jaccard, Morisita, and Tversky) were used to assess the CDR3 repertoire in bats, humans, and mice (Figure 5A). The analysis showed that the bat CDR3 repertoires was variable and varied significantly, with some overlap coefficients slightly higher than humans, but significantly lower than mice.

**Figure5.**
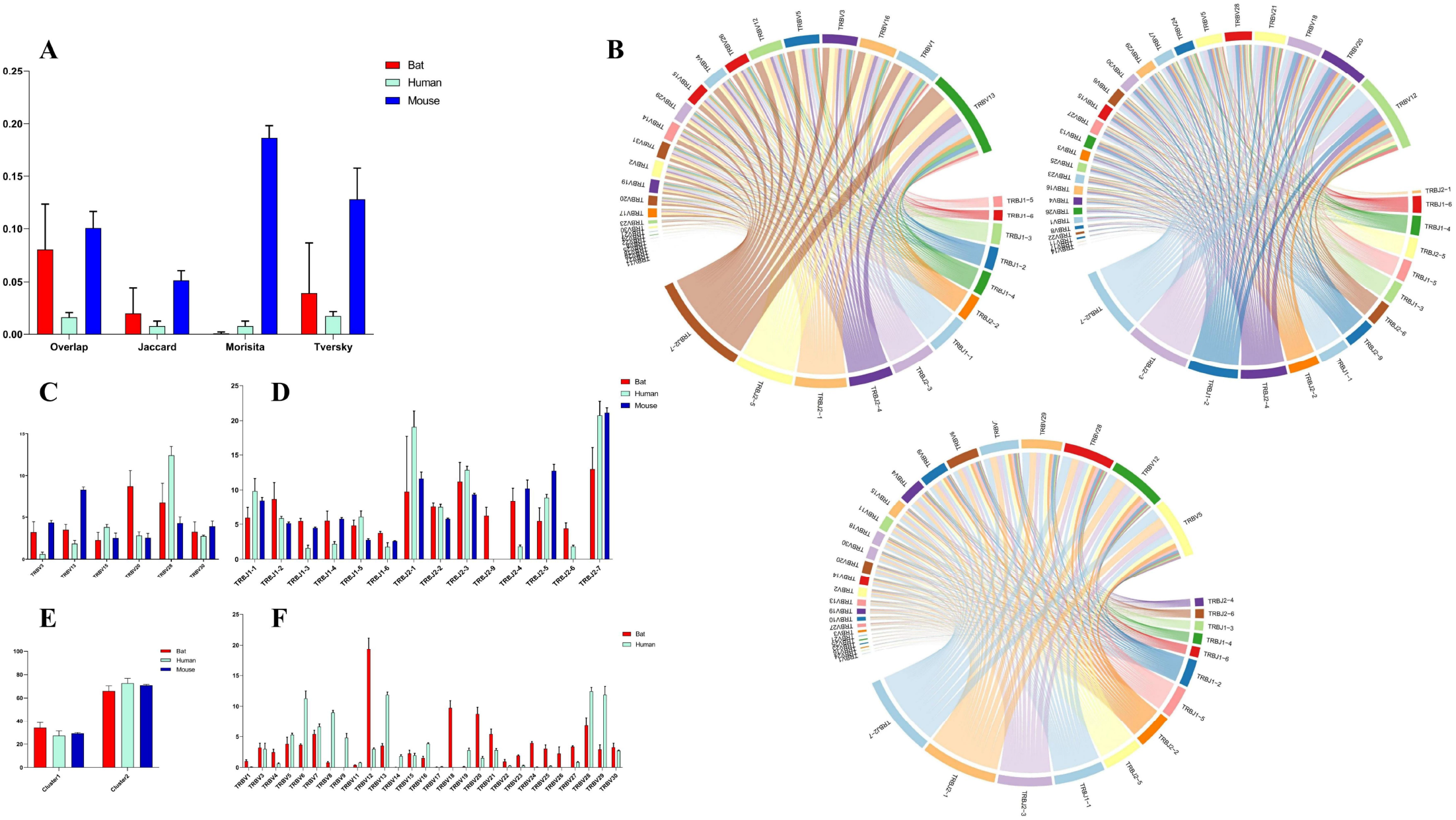
Comparative Analysis of TCR β Chain Repertoire Genes and Motif Usage in Bats, Humans, and Mice. (A) Overlap index of CDR3 sequences in the three species. (B) V-J pairing analysis of the three species, with the order being Batl, Human 1, and Mouse 1. (C) Homologous TRBV gene usage analysis of the three species. (D) TRBJ gene usage analysis. (E) J gene usage in cluster. (F) TRBV gene usage analysis of bats and humans.

Based on the phylogenetic tree result, we analyzed the homology TRBV gene usage and V-J pairing (Figure 5B, Supplementary Figure 5) in bats, humans, and mice. There are six homology TRBV genes expressed, with TRBV15 and TRBV30 exhibiting similar usage (Figure 5C). Furthermore, we compared the usage of all TRBV genes in bats and humans, as shown in Figure 5F. The most commonly used TRBV genes in bats were TRBV12, TRBV18, and TRBV20, whereas in humans, they were TRBV13, TRBV28, and TRBV29. Notably, TRBV9 and TRBV19 were either not expressed or exhibited low expression in the bat samples. Additionally, the TRBV9 group was exclusive to P. discolor, with all four TRBV9 members being pseudogenes. Among the three species, all functional TRBJ genes, with the exception of TRBJ2-8 (pseudogene) in bats and TRBJ2-6 (pseudogene) in mice, were expressed (Figure 5D). Through comparative TRBJ analysis, we observed that high usage of TRBJ2-1, TRBJ2-3, and TRBJ2-7 occurred concurrently in bats, humans, and mice. Additionally, we observed that most of the J genes used in the three species were derived from the second D-J-C cluster (Figure 5E). This finding suggests that there may be a correlation between gene usage during rearrangement and distance.

The CDR3 region motifs (5 amino acids) were analyzed and four of the top ten high-frequency motifs were shared among bats, humans, and mice, with most being CASSN motifs (Figure 6A, Figure 6B). This suggests that the cell populations involved in the immune response in the three species may be similar. The CDR3 region length was also characterized, with a bell-shaped distribution of 14 amino acids for bats and mice, and 15 amino acids for humans (Figure 6C). The length effects caused by insertion and deletion of nucleotides in the CDR3 region were assessed, and it was found that humans and mice had similar deletions at the V’3 end. Bat and mouse had similar insertions at the V 3’ end and the J gene 5’ end (Figure 6D, 6E). Finally, the amino acids in the CDR3 region were consistent, with high frequencies observed from S, G, and A in all three species (Figure 6F).

**Figure 6.**
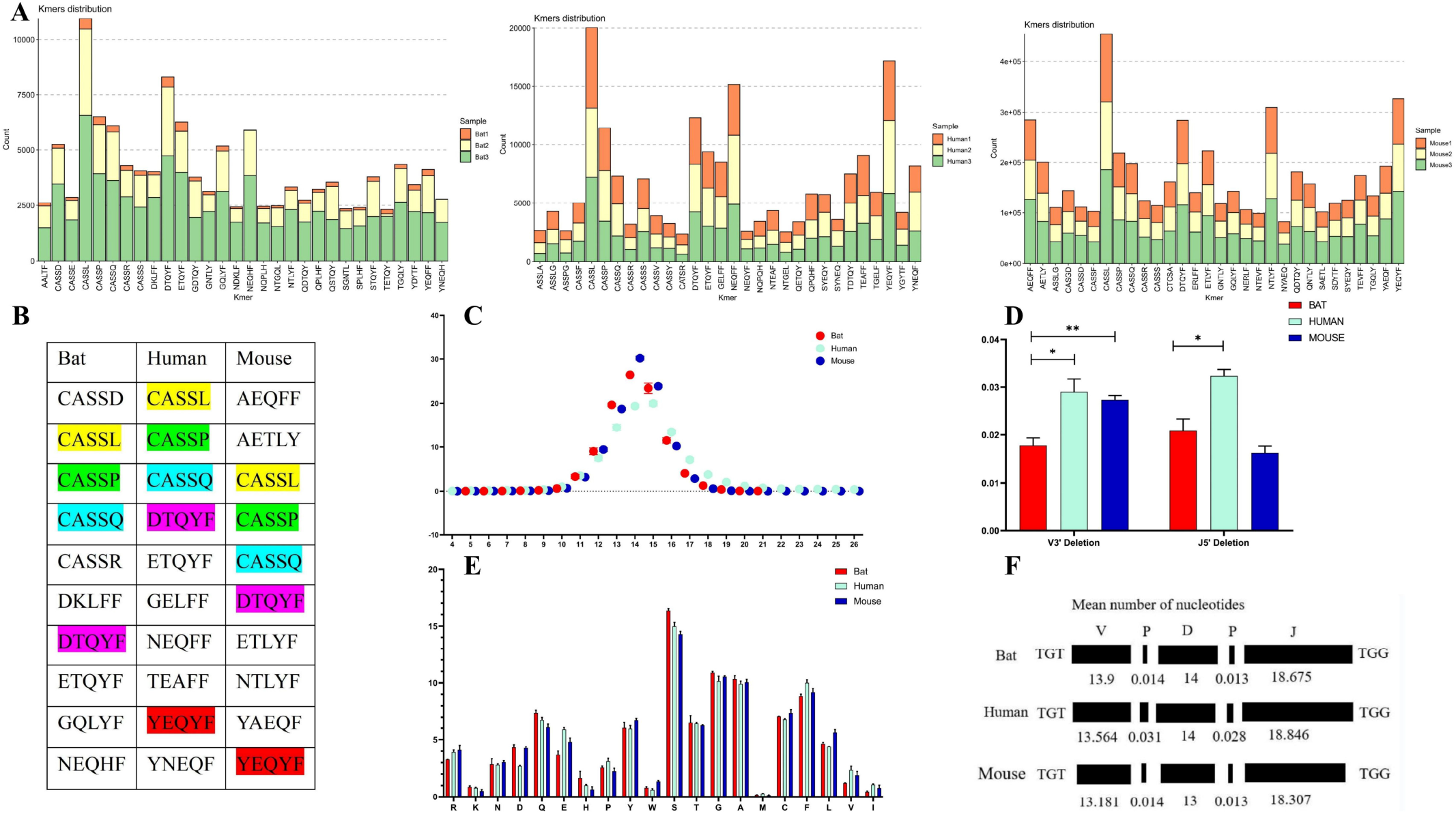
Comparative analysis of the CDR3 region features of the TCRβ chain repertoire in bat, human, and mouse is presented in this figure. (A) The top 30 5 AA length motifs used in the CDR3 region are analyzed. (B) The top 10 shared motifs are labeled in the same color. (C) The CDR3 region length is analyzed. (D) Deletions at the 3’ end of the V gene and 5’ end of the J gene are compared. (E) AA usage in the CDR3 region is analyzed. (F) The composition of the CDR3 region is presented.

## 4 Discussion

The lack of basic genetic information has hindered the study of bat immunity, despite their importance in virus transmission and unique physiological traits. Earlier studies have identified T, B, and Macrophage populations in Indian fox bats, with a higher T/B cell ratio in the spleen and lymph nodes compared to mice(*24*). Further studies found that CD4+ T cells dominate the blood lymph and CD8+ T cells dominate the spleen in fox bats(*25*), suggesting a role of T/B cell immunity in bats, but the understanding of TCR is limited.

The TR/IG loci of the Greater horseshoe bat and the Egyptian fruit bat were annotated separately in 2021 (*20, 26*). However, no public database or tool has been used to analyze the TCR/BCR sequences for bats, and the TCR/BCR repertoire characterization of bats has not been reported. To create a bat germline gene database, the priority work was a complete annotation of TR/IG loci. It should be noted that the TR/IG loci annotation is different from normal genome annotation and it requires a high-quality genome because the shortly genes (like D and J genes) and the loci’s large span on the genome. For our research, we chose three bat species with high-quality genomes from the database: R. ferrumequinum, P. discolor, and P. pipistrellus. Creating germline gene databases involves finding all potential genes based on specific structural features, which is the central issue. Due to the short sequence of D (10-15 bp) and J (30-50 bp) genes, we used the RSS sequence search method simultaneously to avoid losing sequences from the homology search. However, non-classical RSS (12/23+/-1 nt) may affect the final result, but such genes are rare in humans and mice. Moreover, pseudogenes that lack RSS cannot be involved in the rearrangement process, meaning that they have no impact on the productive repertoire and pathogenic response(*27*).

In Chiroptera, previous studies have found varying degrees of replication of immune-related genes, but for TCR, gene duplication events were mostly reported in Artiodactyla such as *cattle(28, 29)* and goats(*30*), which have diverse T cell receptor pools. Surprisingly, massive gene duplication events of the TRBV12 family in P. discolor and considerable variation in TRBV gene numbers among Chiroptera have never been reported in mammals. In comparison, studies of primates(*31–33*), carnivores(*34–36*), and camels(*37*), have shown minimal differences in TRB loci structure and the number of germline genes among species of the same orders. The distribution and evolutionary direction of TRBV gene families in the three bat species is similar to Artiodactyla, with random and massive replications. The lack of gut-associated lymphoid tissue (GALT) in Rhinolophus Hildebrandti and Pipistrellus pipistrellus makes it unlikely that they possess a gene conversion related mechanism to generate receptor diversity as observed in the chicken immunoglobulin system (*38–41*). Instead, T cells use gene rearrangement as the predominant mechanism for generating TCR diversity in bats. This difference in rearrangement likelihood suggests that TCRβ chain repertoire diversity might be present in at least the three annotated bat species. The anchor residues in TRBV genes are important for MHC peptide binding, and while MHC genes in bats are more diverse than in other mammals, previous research has shown high sequence similarity between bat and human, mouse, dog, and cow MHC genes (*42–44*). Moreover, the anchor residues in the annotated TRBV and TRBJ genes of bats are highly conserved, indicating that TCR recognition in bats is MHC-restricted.

We established a bat TCR β chain reference library and analyzed the TCR β chain repertoire using 3 R. affinis samples. From a total of 4.2 million reads on average, we found that the number of TCR clonetypes differed among the bat samples, with bat1 having 3498, while bat2 and bat3 had 25443 and 45771, respectively. There are several possible explanations: Firstly, this bat sample may have little expression of TCRs. It implies that the diversity of individual bats may vary. Second, since this is the first time that we performed repertoire building as well as sequencing of bats, the sequencing depth may have caused an impact, and increasing the number of sequenced samples would help to solve the potential issues of individual bat diversity and low TCR expression due to the limited collection of samples. After analyzing the sequence output of each bat sample, we discovered that 25 TRBV genes and 14 TRBJ gene families were present in every experimental bat. This finding indicates that we have, for the first time, attained achieved a high level of completeness in identifying the TCR β chain CDR3 in bats. Our results provide a valuable technical resource and data analysis tools to assist in the design and optimization of primers for HTS sequencing analysis of TCR repertoire from bats belonging to different families using 5’RACE. The TCR β chain sequences of R. affinis were highly similar to the annotated genes of R. ferrumequinum. Specifically, 99% of the C genes belonged to R. ferrumequinum, and the TRBV and TRBJ genes were nearly identical to the annotated V and J genes of R. ferrumequinum. This suggests that certain immune response genes are genetically conserved among bat species of the same family.

We conducted an analysis on the usage and pairing patterns of TRBV and TRBJ genes in bats, humans, and mice. Despite humans having over 50 and mice over 30 germline V genes, their specific immune response may rely on only one or a few of them(*45*). The low expression levels of several gene families may be explained by a variety of factors. Firstly, it’s possible that the low expression levels are due to R. affinis itself. Additionally, the TRBV17 family has very low homology in the three annotated bats, which suggests that it may undergo significant mutations in the R. affinis genome and therefore not be detected. These factors could contribute to the overall low expression levels of certain gene families. A kind of mutation might lead to functional genes becoming pseudogenes, a situation that is very common across species. For example, the TRBV1 gene in bats is functional gene, while TRBV1 in humans is pseudogene. There are also cases where functional genes are not detected as expressed, for example, TRBV18 is barely expressed in humans but relatively highly expressed in bats; similar to TRBV23, TRBV24, TRBV26, etc. TRBV30, which is located in a specific position downstream of the second C gene and in a position opposite to transcription in bats, humans, and mice (TRBV31), was observed to be common in all three species. Several studies have reported that V and J gene usage during rearrangement correlates with position in the locus (*46–48*). Interestingly, in all three species, over 60% of the J gene is from the second D-J-C cluster. TRBV12-5 in humans and TRBV13-2 in mice are often linked to autoimmune diseases and account for over 50% of NK-T cells. Additionally, mouse MAIT cells express a TCR-α chain with TRAV1 and TRAJ33 and paired β chains of TRBV13 and TRBV19 (*49–53*). Notably, both TRBV12 in bats and TRBV13 in mice are highly utilized V genes. Bats exhibit significant differences from humans in their usage of multiple TRBV genes, indicating a specific biased amplification in the evolution of bat TCR immune response genes. This may be linked to evolutionary pressure or coevolution with viruses.

The usage of TRB CDR3 region amino acids in bats, humans, and mice is highly similar, with a consistent pattern of high-frequency motifs. However, there are more significant differences in motif usage throughout the CDR3 region, indicating that the V-terminal and J-front ends of the CDR3 regions in these mammals are more conserved, and that the typical antigenic selection encountered in evolution may play a role. The length of the TRB CDR3 region is shorter in bats and mice compared to humans, which we attribute to differences in insertion and deletion at the V’3 and J5’ ends across the three species, as well as differences in the evolutionary length of the D gene and V/J involvement in the CDR3 region genes. While the overlap between individual bat CDR3 regions is higher than that of humans, it is significantly lower than that of mice, suggesting a higher concordance of CDR3 regions between different individual BALB/c mice with the same genetic background.

We annotated the complete TRB loci in three species of bats and constructed a database for analyzing the bat TCR repertoire. This allowed us to study the TCR evolution and specific immune response mechanism in bats and provide a new theory, technical tools, and data for comparative analysis of the mechanism of virus tolerance in bats.

## Supporting information

suptable1

suptable2

suptable3

suptable4

suptable5

suptable6

## 5 Acknowledgements

We would like to express our gratitude to Jiang Zhou and Xingliang Wang from Guizhou Normal University for assisting in collecting the bat samples. We also thank ImmuQuad Company for their help in constructing the libraries and conducting high-throughput sequencing of the bat samples. Additionally, we would like to thank the Vertebrate Genome Project workers for their contribution in sequencing the bat genomes and sharing their data. Lastly, we extend our thanks to Dianita S. Saputri for revising the manuscript.

## 6 Author Contributions

Xinsheng Yao, Long Ma and Hao Zhou designed the experiment and wrote the paper, while Long Ma and Hao Zhou conducted the experiments, analyzed data, and created graphs. Other contributors, including Jun Li, Longyu Liu, Dewei Zhou, Yingjie Wu, Xingliang Wang, Jiang Zhou, Qingqing, Ma, and Daron M. Standley assisted with sample collection, data analysis, manuscript revision and library construction for humans and mice.

There is no conflict of interest in this paper, and the experimental sample collection of bats and humans was approved by the ethics committee of Zunyi Medical University, and the informed consent of volunteers was obtained for the collection of human peripheral blood samples, and all sample collection and experiments were performed in strict accordance with biosafety requirements.

## 7 Funding

This study was supported by the National Natural Science Foundation of China (31860257&81860300) and the Guizhou Provincial Hundred Talent Fund [No. (2018) 5637].

## 9 Supplementary Material

Supplementary Data 1 Genbank files for Phyllostomus discolor, Rhinolophus ferrumequinum, and Pipistrellus pipistrellus.

## 10 Data Availability Statement

The datasets of bat repertoire for this study can be found in the Sequence Read Archive (SRA) [https://www.ncbi.nlm.nih.gov/bioproject/PRJNA877449].

## Notes

### Competing Interest Statement

The authors have declared no competing interest.

### Summary of Updates

Manuscript has been re-written.

